# Genetic and Epigenetic Reprogramming of Transposable Elements Drives ecDNA-Mediated Metastatic Prostate Cancer

**DOI:** 10.1101/2025.08.08.668693

**Authors:** Lisanne Mout, Thaidy Moreno-Rodriguez, Giacomo Grillo, Ankita Nand, Tina Keshavarzian, Shalini Bahl, Komaldeep Kang, Matthew Bootsma, Emma Minnee, Stanley Zhou, Kathleen H. Burns, Eva Corey, Peter Nelson, Scott M. Dehm, Shuang G. Zhao, Wilbert Zwart, Felix Feng, David Quigley, Mathieu Lupien

## Abstract

Extrachromosomal DNAs (ecDNAs), which replicate and segregate in a non-Mendelian manner, serve as vectors for accelerated tumor evolution. By integrating chromatin accessibility, whole-genome sequencing, and Hi-C-based genome topology data from a cohort of metastatic Castration-Resistant Prostate Cancer (mCRPC) cases, we show that epigenetically activated repeat DNA, amplified in ecDNAs, drive oncogene overexpression. Specifically, we identify a subgroup of mCRPCs (20%) characterized by clusters of accessible LINE1 repeat DNA elements flanking the androgen receptor (AR) gene. These LINE1 elements are co-amplified with AR and provide binding sites for prostate-lineage transcription factors, including AR, FOXA1 and HOXB13. Accessible LINE1 elements establish novel 3D chromatin interactions with the AR gene, forging a new regulatory plexus driving AR overexpression and confers resistance to androgen signaling inhibitors. Our findings indicate how tumor evolution is driven by the convergence of genetic and epigenetic alterations on repeat DNA, activating and amplifying them to allow oncogene overexpression.

**Statement of significance:** We show how tumor evolution is driven by the convergence of genetic and epigenetic alterations on repeat DNA elements, resulting in their activation as regulatory elements and co-amplification in ecDNAs with oncogenes in mCRPC.

## Introduction

Unlike localized disease, metastatic castration-resistant prostate cancer (mCRPC) is an aggressive disease lacking curative treatments. Consequently, prostate cancer (PCa) is one of the leading causes of cancer related death in men ^1^. Blocking androgen receptor (AR) signaling is the golden standard for managing metastatic PCa but inevitably fails because of acquired resistance mechanisms including reactivation of the AR pathway and epigenetic reprogramming driving neuroendocrine signaling ^2^. The search for alternative treatment options against mCRPC has relied on identifying genetic subgroups in metastatic tumors, including cases of copy number gains and ecDNAs-positivity at the *AR* locus, activating structural variants at the *ERG* locus, and tandem duplication at the *MYC* locus found in ∼80%, ∼40% and 30% of tumors, respectively ^3–5^. Similarly, a DNA methylation-based non-genetic subgroup associated with mutations in the *TET2*, *DNMT3B*, *IDH1*, and *BRAF* genes was reported across ∼20% of patients ^6^. However, only subsets of prostate cancer patients benefit from these discoveries with advanced therapeutics ^7–9^, highlighting the need to identify and characterize the molecular processes active across the mCRPC subtypes to guide the development of new therapies and select treatments tailored to each patient.

Epigenetic characterization of the chromatin landscape in both healthy tissues and cancer effectively identifies phenotypically distinct cell types and states using differentially accessible DNA sequences, also termed chromatin variants ^10–12^. Chromatin variants mark cell type or state-specific cis-regulatory elements located within the non-coding genome, including repetitive DNA sequences, such as transposable elements ^13^. Transposable elements can play many different roles in cancer, from triggering a viral mimicry response by generating double-stranded RNA to introducing somatic mutations through their ability to retrotranspose ^14–20^. More commonly, transposable elements are targets of chromatin variants that lead to their onco-exaptation for gene regulatory activities. Onco-exaptation includes the use of transposable elements as transcription start sites near oncogenes to generate fusion transcripts ^21,22^. Transposable elements can also be co-opted as distal regulatory elements to support oncogene overexpression and to sustain stemness properties, in primary prostate tumors as well as in other cancers ^23–26^.

In the present study, we characterized the genetic and epigenetic landscape and defined phenotypically distinct mCRPC subgroups from chromatin variants marking transposable elements. We identified a LINE1+ subgroup associated with LINE1 transposable elements co-amplified with the *AR* gene that are co-opted from non-genetic variants as regulatory elements to drive *AR* overexpression and response to AR signaling inhibitors.

## Results

### Chromatin variants over transposable elements stratifies mCRPC into four subgroups

To assess whether the chromatin landscape of transposable elements identifies phenotypically distinct mCRPC patients, we assessed their enrichment in accessible chromatin from ATAC-seq data generated across the WCDT cohort (71 samples, from 66 mCRPC patients) ^27^. Unsupervised clustering of transposable element subfamilies (N=605) in accessible chromatin revealed four mCRPC subgroups (Figure 1A), with 196 subfamilies differentially enriched across the subgroups (Supplementary Figure 1A, Supplementary Table 1). The epigenetic subgroups identified from the repetitive genome were independent of previously identified subtypes defined by methylation or genome wide topology data (Supplementary Figure 1A, heatmap annotation).

**Figure 1:**
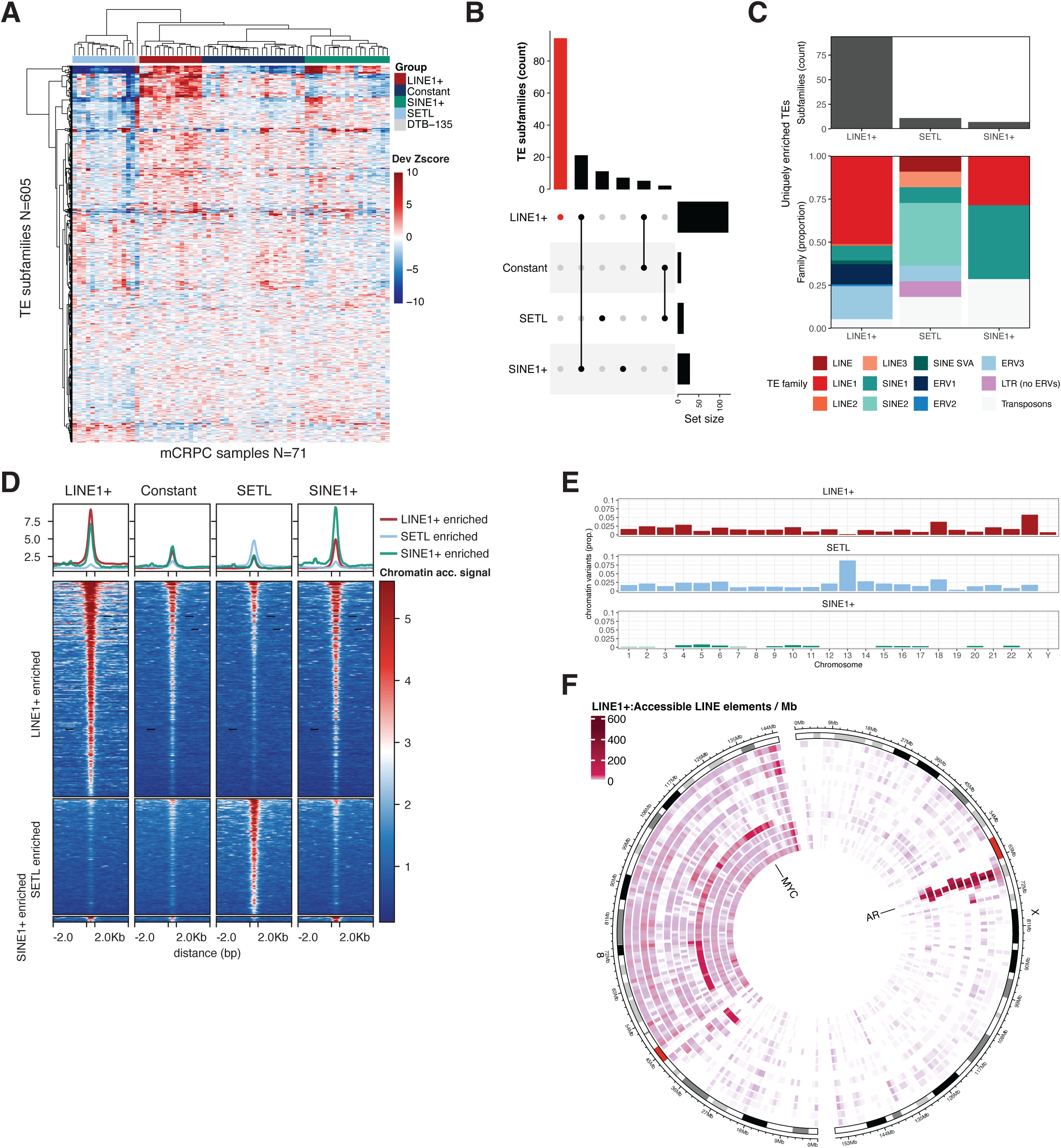
Enrichment of transposable elements define mCRPC subtypes and identify clusters of accessible LINE1 elements at the AR locus. **A)** Heatmap with unsupervised clustering of deviation Z-scores (Dev Zscore) for transposable element subfamilies (N=605, rows) in accessible chromatin from mCRPC samples (N=71, columns). Transposable element subgroups were identified from unsupervised clustering of transposable element enrichment patterns. DTB-135 clustered separately from the four transposable element subgroups identified. **B)** Upset plot displaying the number of enriched transposable element subfamilies (q<0.01; median Z-score >1) shared across the different mCRPC groups. **C)** Barplot showing the transposable element subfamilies uniquely enriched (q<0.01; median Z-score >1) in the different mCRPC groups (top) and allocation to transposable element families (bottom). No transposable element subfamilies were uniquely enriched in the Constant subgroup. **D)** Tornado plot showing the average fold enriched chromatin accessibility signal in the mCRPC groups across differentially accessible elements (q<0.01) from enriched transposable element subfamilies. The length of the elements was uniformly scaled to 500 bp with 2 Kb flanking regions. **E)** Barplot showing the genomic distribution of differentially accessible elements by chromosome, normalized to the number of peaks across transposable elements. **F)** Circos plot showing the density of accessible LINE1 elements from enriched subfamilies in 1 Mb bins for chr8 and chrX. Each track represents a single sample in the LINE1+ group. The positions of the *AR* and *MYC* oncogene are annotated.

The WCDT cohort included five patients with matched pre- and post-treatment samples including three patients who transitioned between transposable element enrichment states upon treatment progression, suggesting that the chromatin state of transposable elements is not static, but instead dynamically altered and reflects distinct phenotypic subtypes (Supplementary Figure 1B). We performed a stringent selection for transposable element subfamilies enriched in accessible chromatin (median Z-score >1) and identified primarily LINEs (N=73), SINEs (N=49), and ERVs (N=37, Figure 1B, Supplementary Table 2, 3). The enrichment of 48 unique LINE1 subfamilies dominated a subgroup of 14 samples, which we labeled “LINE1+” (Figure 1A-C, Supplementary Table 4). Other subgroups were labeled “Constant”, being characterized with no transposable element subfamilies uniquely enriched in accessible chromatin (23 samples), “SETL” showing enrichment across a combination of SINE, ERV3, Transposon and LINE subfamilies (14 samples), and “SINE1+” based on the preferential enrichment of SINE1 subfamilies in accessible chromatin of 19 mCRPC samples (Figure 1A-C).

Most transposable elements can no longer retrotranspose, with the exception of 146 full-length intact elements (FLI-L1) primarily belonging to the L1Hs and L1PA2 subfamilies, which are generally repressed through chromatin compaction ^28,29^. In addition, approximately 100 elements from evolutionary younger LINE1 subfamilies encode a full-length *ORF2* protein, that allows for the retrotransposition of other LINE or SINE elements ^30^. In our analysis the L1Hs and L1PA2 subfamilies were not significantly enriched in accessible chromatin of LINE1+ or other mCRPC samples (Supplementary Table 4). However, up to 22 elements were located in accessible chromatin across all mCRPC subgroups and showed higher levels of chromatin accessibility in LINE1+ (Supplementary Figure 1C). This elevated accessibility reflects an epigenetic context permissive for retrotransposition activity.

To assess if sample and sequencing quality metrics impacted transposable element enrichment and the clustering of samples, we compared biopsy location, peak counts, tumor cell content and fraction read in peaks (FRiP) across the transposable element groups (Supplementary Figure 1D-E, Supplementary Table 5, 6). This identified significantly lower peak counts, FRiP and tumor cell purity in the Constant samples (Supplementary Figure 1E). However, repeating the transposable element-based clustering of samples with FRiP > 0.1 and tumor cell content >50% (N=55) largely replicated our subgroups (Supplementary Figure 1F), showcasing that transposable element enrichment in accessible chromatin was robustly observed. In addition, we identified a similar overrepresentation of transposable element families in accessible chromatin using the cohort of LuCaP patient-derived xenograft (PDX) models (N=39 samples from 38 models, Supplementary Figure 1G-I, Supplementary table 7) ^31^. These findings establish four mCRPC subgroups based on the variability in chromatin accessibility over repetitive DNA sequences.

### Chromatin variants marking LINE1 elements cluster near the AR gene locus

To uncover transposable elements co-opted for their cis-regulatory potential by oncogenic drivers we identified chromatin variants enriched in the different epigenomic mCRPC subgroups ^12^. The number of chromatin variants across the genome varied per subgroup, ranging from 10,250 in the SETL subgroup to 19 in the Constant subgroup (Supplementary Figure 2A-B). We selected chromatin variants with increased chromatin accessibility found at transposable elements unique to the subgroup-specific subfamilies. This identified 27 variants in SINE1+, 560 in SETL and 1,042 in the LINE1+ group that primarily resided in intronic and distal intergenic regions, consistent with their activation as cis-regulatory elements (Supplementary Figure 2A). Chromatin variants enriched in the LINE1+ subgroup showed reduced chromatin accessibility levels across the other subgroups, while repeat elements enriched in the SETL subgroup lacked accessibility in all other subgroups, suggesting that the SETL subgroup is most distinct from all other subgroups (Figure 1A, D).

Next we assessed the distribution of chromatin variants from the different mCRPC subgroups across chromosomes. In LINE1+ mCRPC samples, chromatin variants over transposable elements were found preferentially on the X chromosome where the *AR* gene is located (Figure 1E). Transposable elements are not randomly distributed within the human genome due to nonrandom insertions generated by retrotransposition events, enhanced by natural selection and genetic drift ^28^. Elements from 13 LINE1 subfamilies were found to preferentially occupy the AR amplified region on chrX (Supplementary Figure 2C).

Visualizing the density of accessible transposable elements from LINE1 subfamilies revealed a hotspot at the *AR* gene locus in LINE1+ mCRPCs, in comparison to the density seen across the genome such as at the *MYC* gene locus on chromosome 8 (Figure 1F, Supplementary Figure 2D-G). A total of 68 transposable element subfamilies harbored significant enrichment of accessible elements at the *AR* gene locus in over 50% of the LINE1+ mCRPC samples, the majority (46/68) belong to the LINE1 family (Supplementary Figure 2H, Supplementary Table 8). Clusters of accessible transposable element subfamilies at the *AR* gene locus were rare in SINE1+, SETL, and Constant mCRPC samples. These findings establish LINE1+ mCRPC as a distinct entity marked that preferentially hijacks LINE1 transposable elements that occupy the *AR* gene locus.

### LINE1+ mCRPC subgroup is associated with AR locus high copy number gains

Next we assessed the relationship between the four mCRPC subgroups and the genetic variants found in these tumors. Using matched whole genome sequencing data from 69 of the 71 samples, we detected a significant bias for high copy number gains of *AR* gene in the LINE1+ subgroup as opposed to the other mCRPC subgroups (Figure 2A, Fisher exact test P=0.02). Further inspection of the *AR* locus showed that amplifications in LINE1+ samples converged with clusters of accessible LINE1 elements and increased chromatin accessibility compared to the other subgroups (Supplementary Figure 3A, B). *AR* gene amplification in LINE1+ samples ranged from 12 to 149 copies with a median of 29 (Figure 2B, Supplementary Tables 9). LINE1 status was associated with increased *AR* RNA expression levels compared to SINE1+, Constant and SETL samples (Figure 2B, Supplementary Figure 3C, Supplementary Tables 10). In addition, all LINE1+ samples were classified as AR-driven disease, according to previously established transcriptomic subtypes (Figure 2C) ^27,32^. The majority of AR-negative samples (N=10) were identified in the SETL subgroup, including neuroendocrine (AR-NE+, N=3) and double negative samples (AR-NE-, N=4). Loss of AR signaling has been attributed to epigenetic reprogramming which is in line with our previous observations of SETL samples having a distinct chromatin landscape ^33^. To validate the relationship between *AR* expression and the LINE1+ phenotype, we scored LuCaP PDX models using a signature composed of LINE1 subfamilies uniquely enriched in the LINE1+ mCRPC samples (Supplementary Figure 3D). LuCaP models that scored within the top 33% for the LINE1 signature, showed increased AR expression (P=0.002) and pathway activity (P=0.06) with decreased NE pathway activity as compared to the lowest 33% (P=0.04, Supplementary Figure 3E). Overall, these findings highlight a strong association between LINE1+ mCRPC and *AR* gene amplification, suggesting that transposable element accessibility may contribute to *AR* overexpression and reinforce the AR-driven nature of this distinct tumor subgroup.

**Figure 2:**
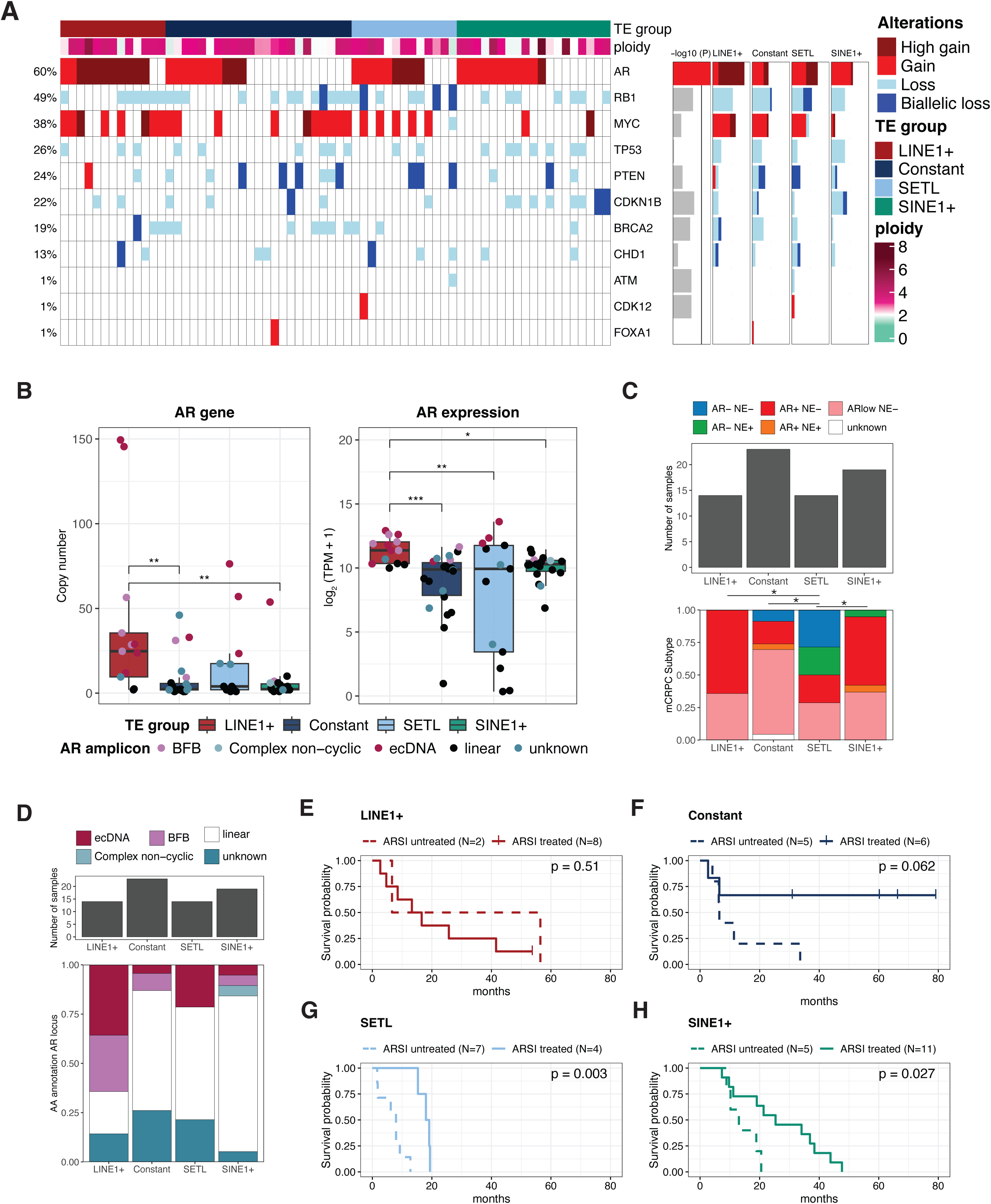
Enrichment of accessible LINE1 elements is associated with recurrent AR amplifications in mCRPC. **A)** Oncoprint showing oncogenic drivers with recurring copy number alterations across 69 mCRPC samples with matched ATAC-seq and WGS. For *AR*, weighted copy numbers exceeding 0.9*ploidy were annotated as gains and >5*ploidy as high gains. For genes on autosomes weighted copy numbers exceeding 1.95*ploidy and 3*ploidy were annotated as gains and high gains respectively. Losses refer to a weighted copy number below 1.1 and biallelic loss below 0.5*ploidy. The barplots (right) summarize the results of the statistical comparisons (Fisher exact test, line represents P=0.05) and proportion of recurring copy number alterations across the transposable element subgroups. **B)** AR weighted copy number (left) and normalized expression transcripts per million (TPM; right) across the different transposable element subgroups annotated for the identified amplicon type. Statistical comparison was performed by a Kruskal-Wallis test with Dunn post-hoc test and Benjamini-Hochberg corrections. For *AR* copy-number, LINE1+ vs Constant and SINE1+ P=0.006, P=0.008 respectively. For *AR* expression, LINE1+ vs Constant, SINE1+ and SETL, P=7*10^-4^ and P=0.0047 and 0.0084 respectively. **C)** Transcriptional subtypes based on AR and neuroendocrine signaling across transposable elements subgroups. Statistical comparison was performed by a Fisher exact test grouping AR+NE- and ARL_NE, followed by pairwise comparisons with Benjamini-Hochberg corrections, LINE1+, Constant and SINE1+ vs SETL P=0.02, P=0.03 and P=0.02 respectively. **D)** Barplot showing the proportion of AR ecDNA (N=11), BFB (N=7) and complex non-cyclic alterations (N=1) across the different transposable element groups as determined by AmpliconArchitect. Cases where no complex rearrangement was identified were annotated as linear. Statistical comparison of AR ecDNA and BFB amplicons between the different transposable element groups was performed using a Fisher exact test, with P=0.03 for ecDNA and P=0.07 for BFB. **E-H)** Overall survival with respect to ARSI treatment status across transposable element subgroups in the WCDT cohort (N=48). Solid lines represent ARSI treated patients and the dotted lines represent untreated patients. Statistical comparison was performed using the log-rank test.

High copy number gains and oncogene expression levels are reported in tumors with extrachromosomal DNA (ecDNA), a circular amplicon that spans up to several Mbp’s with low nucleosome density that allows for the activation of regulatory DNA elements ^34,35^. Hence, we used the AmpliconArchitect tool on the whole-genome sequencing data to assess the presence of ecDNAs and related copy number alterations in the mCRPC patients. Recurring *AR* gene locus ecDNA and breakage fusion bridge (BFB) amplicons, which generated similar copy number gains to ecDNA amplifications were found in 16% (N=11) and 10% (N=7) of the mCRPC samples, respectively (Figure 2D, Supplementary Figure 3D, Supplementary Table 11). The proportion of ecDNA and BFB positive samples across the four mCRPC subgroups was significantly different (Fisher test, P=0.03 and P=0.07 respectively). Of note, LuCaP 105CR scored within the highest ranking models for the LINE1 signatures, harbors an AR ecDNA amplification ^36^. Hence, LINE1+ mCRPC is associated with *AR* ecDNA and BFB amplifications, driving high copy number gains, *AR* overexpression, and enrichment of co-amplified LINE1 elements in accessible chromatin.

Resistance to AR-signaling inhibitors (ARSI) is often considered to arise from increased AR pathway activity ^37^. We therefore investigated if LINE1+ mCRPCs were intrinsically refractory to ARSI therapy. Using survival data in the context of next-line ARSI treatment collected on 48 of our 66 mCRPC patients, we detected improved outcomes in SINE1+ and Constant mCRPC patients from ARSI treatment (P=0.03 and P=0.06), with ∼70% of Constant mCRPC subgroup patients presenting a long-term survival with ARSI treatment of over 80 months (Figure 2E-H, Supplementary Table 12). No significant survival benefit was observed in LINE1+ mCRPC patients treated with ARSI (P=0.51). The SETL subgroup benefited from ARSI treatment (P=0.003), although the effect was short-lived with all patients dying within 20 months (Figure 2F, Supplementary Figure 3F). Recognizing the small cohort size, these results suggest that LINE1+ mCRPCs exhibit limited benefit from ARSI therapy, and warrant further investigation to support a role for transposable elements in driving an AR hyperactive phenotype of relevance to treatment selection.

### Accessible LINE1 elements are cis-regulatory elements for prostate cancer transcription factors

We previously reported the role of transposable elements as regulatory elements to favor signaling through the AR transcription factor in localized primary prostate cancer ^23^. The clustering of accessible transposable elements at the *AR* gene locus in the LINE1+ mCRPC led us to assess if these could be acquired regulatory elements to enhance *AR* gene expression. Using the ReMap ChIP-seq database, we compared the chromatin binding profiles (termed cistromes) of 1,210 transcription factors generated across 737 different cell states against the accessible transposable elements from 48 LINE1 subfamilies enriched in LINE1+ samples. This identified recurrent enrichment for prostate cancer-related transcription factors, specifically AR, its cofactors FOXA1, HOXB13, GATA2, and co-repressor TLE3 over 19 to 34 LINE1 subfamilies (Figure 3A, Supplementary Table 13)^38–41^. In contrast, accessible transposable elements enriched in LINE1+ mCRPC from the SINE, ERV, and Transposon subfamilies (N=46) were associated with the binding of chromatin remodelers including TRIM28 (Supplementary Figure 4A, Supplementary Table 14) ^42^. The AR cistrome was also enriched across 12 transposable element subfamilies, mainly ERVs, albeit with lower enrichment scores compared to TRIM28 (Supplementary Figure 4A-B). The accessible LINE1 transposable element subfamilies most strongly linked to AR, FOXA1, HOXB13, GATA2 and TLE3 binding in prostate cancer were the mammalian LINE1 subfamilies and L1PB1 (Figure 3B-D, Supplementary Table 14).

**Figure 3:**
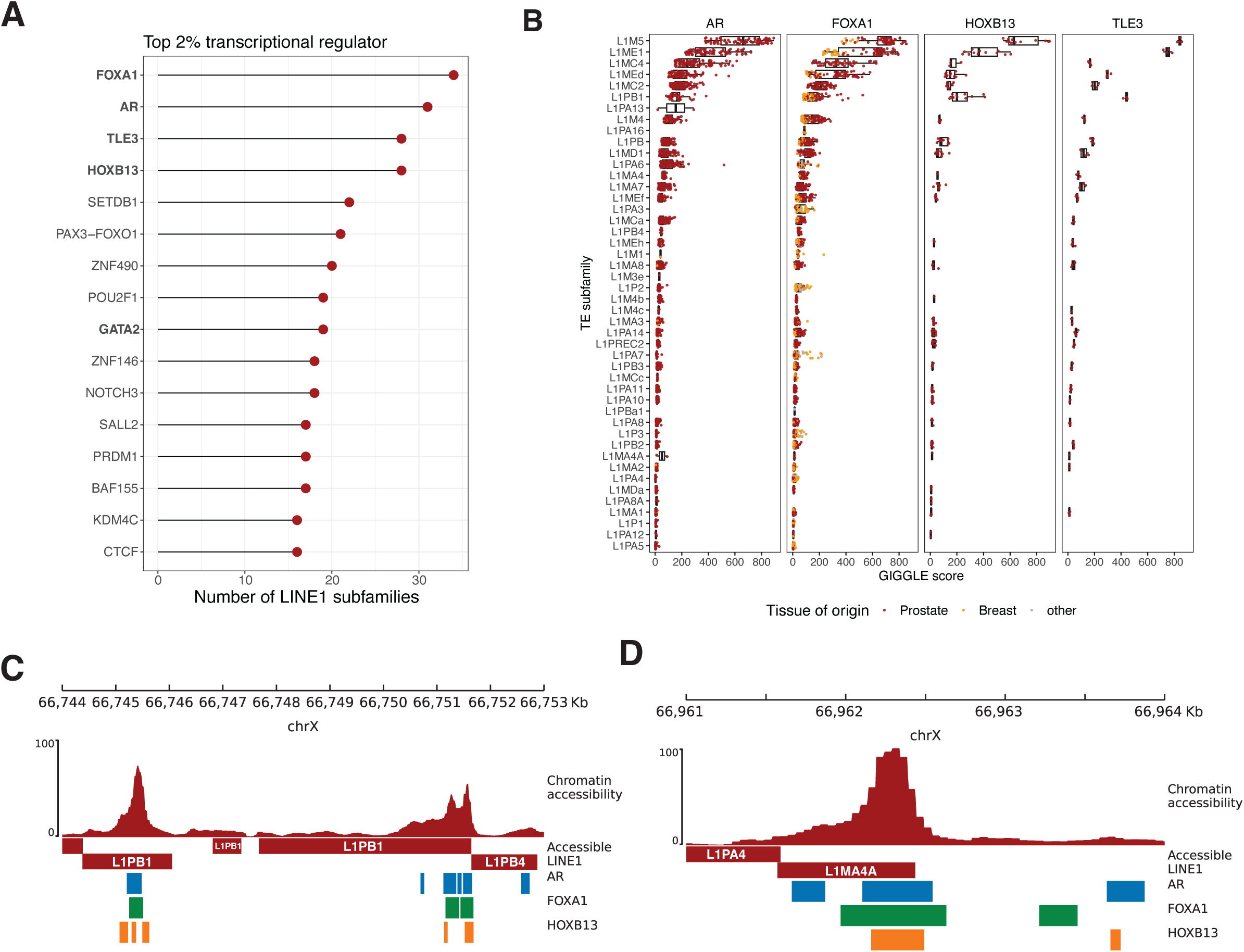
Accessible LINE1 elements provide AR, FOXA1, and HOXB13 binding sites in mCRPC. **A)** Top 2% recurring transcriptional regulators with significant enrichment (q=0.05, log-odds ratio >1.5) over LINE1 subfamilies (N=46) as identified from ReMaP. **B)** GIGGLE scores for top transcriptional regulators over enriched LINE1 subfamilies identified from the ReMaP analysis. **C-D)** Case example of accessible L1PB1 **(C)** and L1MA4A elements **(D)** in LINE1+ samples with transcription factor binding sites as identified from ReMaP. The top track depicts the average fold-enriched chromatin accessibility signal, with the second track showing accessible LINE1 elements and AR (blue), FOXA1 (green), and HOXB13 (orange) binding sites.

These observations suggest that accessible LINE1 transposable elements co-amplified with the *AR* gene serve as acquired regulatory elements in LINE1+ mCRPC, co-opted by AR and its cofactors.

### Accessible LINE1 elements define a new regulatory plexus driving AR overexpression

To identify whether accessible LINE1 transposable elements regulate *AR* gene expression in LINE1+ mCRPC, we integrated the chromatin accessibility landscape with genome topology generated from matched HiC data. In line with previous reports linking HiC signal spanning the *AR* enhancer to gene locus (644 Kb) with ecDNA and BFB status ^43^, we found significantly lower regional contact frequency sliding window (RCFS) scores in LINE1+ samples compared to Constant and SINE1+ samples (P=0.01 and P = 0.009 respectively, Figure 4A, Supplementary Figure 5A Supplementary Table 16). This was most pronounced in the ecDNA and BFB-positive samples (Supplementary Figure 5A). Restricting the HiC data analysis to chromatin loop detection revealed an increased number of contact interactions (loops) linking the *AR* transcription start site (TSS) and distal accessible regions within the amplified *AR* locus. We identified a median of 424 interactions in LINE1+ samples compared to 40, 69, and 91 in the Constant, SETL and SINE1+ groups respectively (P=0.005, P=0.009 and P=0.05 respectively, Figure 4A). This suggests that LINE1+ samples reshape their chromatin topology to establish a new regulatory plexus for the *AR* gene.

**Figure 4:**
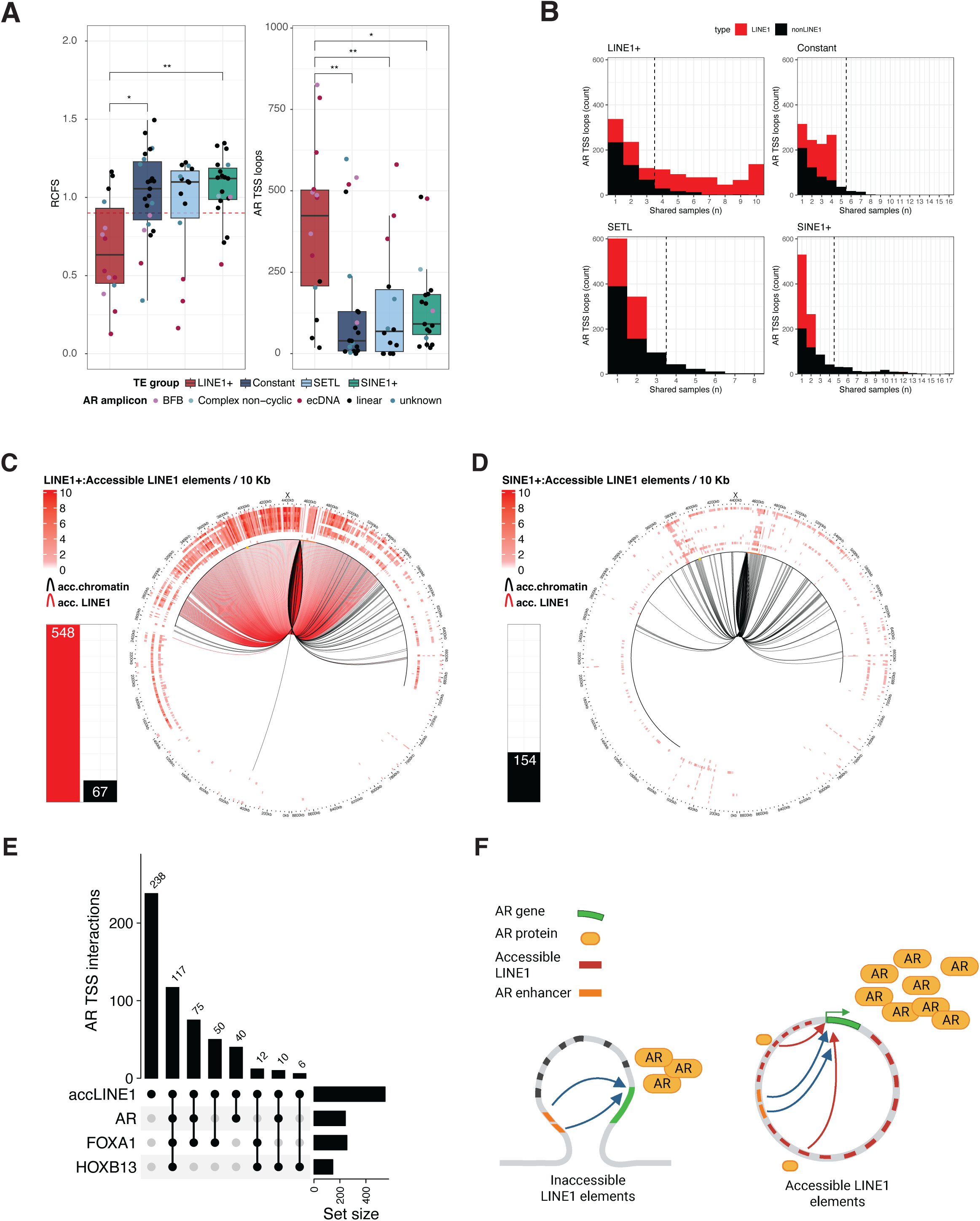
Accessible LINE elements are cis-regulatory elements that interact with the *AR* TSS. **A)** Mean scaled regional contact frequency score (RCFS) for the *AR* amplified gene locus, spanning the transcription start site (TSS) to canonical enhancers, across the transposable element subgroups (left). The red dotted line indicates the RCFS cut-off (0.9) associated with AR ecDNA status^43^. AR TSS (+\- 10Kb) intrachromosomal loops (right) anchored to regions containing accessible DNA elements (chrX:65-70Mb). Statistical comparison was performed using a Kruskal-Wallis test with Dunn post-hoc test and Benjamini-Hochberg corrections. RCFS in LINE1+ vs Constant and SINE1+ P=0.01 and P=0.009 respectively. TSS loops in LINE1+ vs Constant, SETL, and SINE1+ P=0.04, P=0.04, and P=0.05 respectively. **B)** *AR* TSS intrachromosomal loops anchored to regions with accessible DNA elements shared across samples. Loops anchored to accessible DNA elements containing LINE1 sequences from enriched subfamilies are shown in red, with the remainder shown in black. The black dotted line represents the cut-off for identifying recurrent interactions in each transposable element group (25%). **C-D)** Circos plots showing recurring (>25%) *AR* TSS intrachromosomal loops in LINE1+ **(C)** and SINE1+ **(D)** samples within chrX:63-71.8Mb (corresponding to cytoband q11.2-q13.1). Loops (inner track) anchored to accessible DNA elements with or without LINE1 sequences from enriched subfamilies are shown in red and black respectively. Barplot summarizes the number of recurrent loops shown in the Circos plot. The *AR* amplified region for the LINE1+ and SINE1+ group is marked with a black line which includes the *AR* gene in orange and the canonical enhancers in yellow. The outer tracks show the density of accessible LINE1 elements from enriched subfamilies in 10 kb bins, for each sample in the subgroup. **E)** Upset plot showing recurrent *AR* TSS loops (>25%) to accessible LINE1 sequences from enriched subfamilies identified in LINE1+ samples overlapping AR, FOXA1 and HOXB13 binding sites derived from ReMaP. **F)** Schematic overview of the proposed model where a novel regulatory plexus is established through co-amplification of LINE1 elements with the *AR* gene and co-opted by the AR transcription factor and its cofactors to favor *AR* gene overexpression.

We next evaluated accessible LINE1 elements across the *AR* gene regulatory plexus (Figure 4B). Selecting interactions recurring in ≥25% of the samples, we identified 615 chromatin interactions linked to the *AR* TSS in LINE1+ samples. In total, 89% (N=548) of these chromatin interactions were anchored to regions containing accessible LINE1 elements (Figure 4C). Recurring chromatin interactions with the *AR* TSS shared in Constant, SETL, or SINE1+ samples were not connected to accessible LINE1 elements (Figure 4D, Supplementary Figure 5B-C) and included loops with a previously reported *AR* gene enhancer ^44^. Our results suggest LINE1 elements are hijacked as cis-regulatory elements in the *AR* regulatory plexus of LINE1+ mCRPCs.

To identify whether accessible LINE1 elements bound by AR, FOXA1, and HOXB13 provide distal regulatory elements to the *AR* and establish a positive feedback loop, we intersected recurrent *AR* TSS interactions with binding sites provided by the ReMap database. Close to 57% of the *AR* TSS interactions with accessible LINE1 elements overlapped with transcription factor binding sites with co-binding of AR, FOXA1, and HOXB13 found in 117 interactions (Figure 4E). Visualizing the *AR* TSS interactions coinciding with accessible LINE1 elements bound by AR, FOXA,1 and HOXB13 showcased that the majority resides within the *AR* gene amplified locus (Supplementary Figure 5D-F).

Overall, we report a novel regulatory plexus for the *AR* gene in LINE1+ mCRPC. This regulatory plexus is composed of accessible LINE1 elements co-amplified with the *AR* gene and co-opted by the AR transcription factor and its cofactors to favor *AR* gene overexpression (Figure 4F).

## Discussion

Our study employed genome-wide profiles of chromatin accessibility from ATAC-seq assay to identify four epigenomic mCRPC subgroups based on chromatin variants over repetitive DNA sequences. These subgroups displayed unique phenotypic properties including differential response to AR targeted therapy. While we detect *AR* gene locus high copy number gains more frequently in the LINE1+ mCRPC subgroup, patient tumor stratification from genetic variants does not fully reflect the epigenomic subgrouping ^44^. Our subgrouping also differs from the classification of mCRPC from the epigenomic signal across the non-repetitive genome, which rather reflects the AR-driven or neuroendocrine transcriptomic subtypes ^27^. Instead, the repetitive genome-based epigenomic stratification of mCRPC more closely aligns with the heterogeneity reported from genome topology assessment based on HiC data, identifying a subset of *AR* gene locus amplified samples that have limited benefit from AR targeted therapy ^43^. Hence, our insights underscore the distinct biology of chromatin accessibility across the repetitive genome in shaping mCRPC progression and highlight the necessity of integrating epigenomics with genomics data in clinical practice to refine patient stratification and improve interventions.

We also demonstrate that LINE1 transposable elements are co-amplified with the *AR* gene and engage as regulatory elements to drive its overexpression in LINE1+ mCRPC patients. This observation aligns with the adaptive nature of AR signaling for castration-resistant progression ^45^. Transposable elements from the LINE1, ERV1, SINE1, and Transposons families are co-opted as regulatory elements in localized primary prostate tumors ^23^. However, the cluster of LINE1 elements accessible at the *AR* gene locus and their co-amplification as linear, BFB, or ecDNAs reported in mCRPC is not seen in localized primary prostate tumors. Instead, localized tumors hijack a subset of transposable elements normally engaged as regulatory elements in pluripotent stem cells ^23^. Hence our study showcases that cancer cells can take advantage of the reservoir of cis-regulatory elements provided by different transposable element subfamilies to drive disease progression and treatment resistance.

Here we also report on changes to the chromatin state of LINE1 elements coinciding with amplification of oncogenic drivers, in particular through ecDNA and BFB amplicons. Recent studies reported on the strong selection of ecDNA amplifications in the progression from localized to metastatic PCa and with castration resistance using xenograft models ^36,46^. In addition to generating high copy numbers and expression of cargo oncogenes, ecDNA co-amplifies enhancer sequences that can establish new inter and intrachromosomal enhancer oncogene interactions ^47–49^. ecDNAs containing only regulatory elements have been shown to co-exist alongside ecDNAs harboring cargo-oncogenes and suggested to function as trans-interacting regulatory hubs ^49,50^. Interestingly, LINE1 retrotransposition events have been linked to large-scale genome rearrangements including the generation of BFB amplicons and the formation of micronuclei leading to chromothripsis events in model systems ^19,51^. Chromothripsis is a key driver of the generation of ecDNA amplicons, although LINE1 insertions at ecDNA breakpoints have yet to be reported ^52,53^. Similarly, micronuclei cycling is linked to chromatin variant genesis ^54,55^. Whether LINE1 elements are involved in ecDNA-driven genomic alterations and/or increased chromatin variation underscores a critical, yet underexplored, aspect of prostate cancer evolution.

Altogether, our findings highlight the interplay between genetic and epigenetic alterations affecting repetitive DNA sequences as a key determinant of mCRPC classification, positioning LINE1+ mCRPCs as a unique, treatment-resistant subset of tumors and emphasizing the crucial influence of transposable elements in the progression of advanced prostate cancer.

## Methods

### Transposable element enrichment in accessible chromatin

ATAC-seq data from the WCDT cohort of mCRPC samples was generated as previously described ^27^. Data processing was performed using the pipeline available through the CoBE platform (www.pmcobe.ca; ATAC-seq_ML). In short, reads were aligned to the human genome (GRCh38) using Bowtie2 (RRID:SCR_016368), removing duplicate and multi mapped reads and accessible regions (aka: peaks) were called with MACS2 (RRID:SCR_013291) ^56^. Samples with at least 15000 peaks and 10 million unique reads were used in downstream analysis. A consensus catalog of non-redundant peaks was generated using an iterative approach adapted from the ArchR pipeline (RRID:SCR_020982) ^57^. In short, summits provided by MACS2 were extended with 250 bp in both directions, overlapping peaks were sorted by signal intensity and redundant peaks were removed. ENCODE blacklist regions in addition to promoter peaks were removed as they harbor limited cancer and sample specificity ^11,58^.

Transposable element enrichment was performed using ChromVar version 1.28 (RRID:SCR_026570) as described previously ^23,59^. In short, the consensus was used to score the presence or absence of peaks overlapping transposable elements and assigned to 971 subfamilies. ChromVar subsequently calculated bias-corrected deviation scores and Z-scores using randomized background peaks matched in GC content. Transposable element subfamilies for which ChromVar was not able to calculate deviation Z scores in any of the samples were omitted from downstream analysis. Z scores were capped at −10 and 10, used to perform unsupervised clustering of samples using the Ward D2 method and Euclidean distance. Transposable element subgroups were determined using a clustering height cut-off at 150.

Differentially enriched transposable element subfamilies were obtained using a non-parametric Kruskal-Wallis rank sum test with Benjamini Hochberg corrections comparing Z-scores between samples assigned to the different groups (q-value <0.01). To obtain uniquely enriched transposable element subfamilies, we calculated the median Z-scores for the different groups and selected subfamilies with a Z-score ≥1 in one of the subgroups. The results were visualized in an upset plot using the ComplexHeatmap package version 2.22 (RRID:SCR_017270) ^60^. Subfamilies (L1PA4, AluJb, etc.) were assigned to transposable element families (LINE1, SINE1, etc) according to Repbase ^61^ (https://www.girinst.org/repbase/) and dominant transposable element families were used to annotate the different subgroups.

Chromatin variants between samples in the different transposable element subgroups were obtained using Diffbind version 3.8.4 (RRID:SCR_012918)^62^. Significantly enriched chromatin variants for each transposable element subgroup were obtained by combining results from each cross comparison (FDR <0.01, Fold change >1), removing overlapping regions and intersecting with enriched transposable element subfamilies using BEDTools version 2.30.0 (RRID:SCR_006646)^63^. Peak annotation of the chromatin variants was performed using ChIPseeker version 1.42 (RRID:SCR_021322)^64^, with +/− 2 Kb for TSS regions. Chromatin accessibility signal across chromatin variants were visualized using the computeMatrix and plotHeatmap functions from the deepTools suite version 3.5.2 (RRID:SCR_016366)^65^.

The distribution of accessible elements from enriched LINE1 subfamilies was counted for each sample in 1Mb (Figure 1F, Supplementary Figures 2C-F) or 10 Kb (Figure 4C-D, Supplementary Figure 5B-F) windows using Bedtools and visualized using Circlize version 0.4.16 (RRID:SCR_002141)^66^. We performed a permutation test to identify significant enrichment of (accessible) elements from 112 uniquely enriched subfamilies in the *AR* amplified locus (chrX:63800250-69691422) using regioneR version 1.24.0 ^67^. For each sample, the null distribution was generated by randomizing accessible elements 10,000 times throughout the genome, not allowing for overlaps and excluding blacklist regions. Significant deviations from the null distribution (P <0.01) were visualized using complexHeatmap version 2.22.0 ^60^, removing transposable element subfamilies without any significant results across samples.

### Copy number calling and amplicon annotation from WGS

Matched WGS was available from 69 samples and processed using the hmftools suite (https://github.com/hartwigmedical/hmftools) ^44^. Structural variants were determined using GRIDSS version 2.12.2 ^68^, outputs were filtered and post-processed using GRIDSS version 1.11. Read depth ratios and GC normalization were performed by COBALT version 1.11 and B-allele frequency (BAF) was calculated by AMBER version 3.5. Copy numbers were calculated with PURPLE version 3.0, by using outputs from GRIDSS, Strelka, COBALT, and AMBER. Copy number alterations were called using the cut-offs previously determined by Lundberg and colleagues ^32^ with the following modifications; high gains for genes on chromosomes X were called when the minCopyNumber exceeded 5 times the ploidy and high gains on autosomes were called when the minCopyNumber exceeded 3 times the ploidy. Genes with recurrent alterations were visualized in an oncoprint (Figure 2A) and generated using complexHeatmap version 2.22.

Copy number alterations spanning the *AR* gene locus were investigated using AmpliconArchitect (RRID:SCR_023150)^69^ following the authors’ recommended settings with modifications described below. First, fastq files were re-aligned against the AmpliconArchitect curated repository genome reference GRCh38 using a dockerized version of *PrepareAA* version 0.1203.1. Copy number calls generated as described above using PURPLE were smoothed using the *seed_trimmer.py* script from *PrepareAA* with default parameters (--minsize 50000 and --cngain 4.5). The output bed file was provided to the *amplified_intervals.py* script from AmpliconArchitect version 1.2 with optimized parameters (--gain 5 --cnsize_min 100000). The *AmpliconArchitect.py* script was executed with the seed interval list, mapped reads, and parameters (--ref GRCh38 --downsample −1 –extendmode EXPLORE --sensitivems False --plotstyle small --insert_sdevs 3.0 --pair_support_min 2). Classification of the identified amplicon structures was performed by AmpliconClassifier version 0.4.9 with the following parameters: (--ref GRCh38 and --plotstyle individual --min_flow 1 --min_size 5000 --decomposition_strictness 0.1).

Genomic overview of the *AR* amplified locus including chromatin accessibility signal and accessible LINE1 elements from enriched subfamilies was performed using pyGenomeTracks version 3.8 (RRID:SCR_025312)^70^.

### AR expression in the transposable element subgroups

RNA-seq data from the WCDT samples was generated as previously described ^32,44^ and is available through https://quigleylab.ucsf.edu/data. To compare AR expression and pathway activity across the different transposable element subgroups, transcripts per million (TPM) were used. Pathway activity was scored using the Androgen signalling Hallmarks and Beltran neuroendocrine (NE) ^71^ gene sets using the R package singscore, as described previously by Shrestha and colleagues ^27^. Transcriptional subtype annotation, as shown in Supplementary Figure 3D, was previously generated by Shrestha and colleagues^27^.

### AR interactions from HiC data

Regional contact frequency sliding window scores (RCFS), significant interactions for chromosome X and TAD subtypes were obtained from Zhao and colleagues ^43^. Local contact interaction scores for the *AR* gene locus were obtained by mapping mean scaled contact frequency (CF) from the canonical enhancers to the gene locus (chrX:66900400-67545147) using Bedtools version 2.30 and compared for the different transposable element subgroups. To identify loops originating from the *AR* TSS we extended the TSS (chrX: 67545147) by 10 Kb in both directions and counted the number of unique interactions for each sample. We subsequently extracted AR interactions mapping to accessible chromatin by extending the interacting locus with 10Kb in both directions and intersecting with peaks called in the matched sample. We repeated this process for peaks overlapping elements from enriched LINE1 subfamilies and visualized the shared results for each transposable element subgroup using Circlize version 4.16 ^66^.

### ARSI response across the transposable element subgroups

Overall survival data for the WCDT cohort and exposure to ARSI (enzalutamide of abiraterone) was obtained from Zhao and colleagues ^43^. We identified 48 patients allocated to the transposable element subgroups using ATAC-seq with matching survival data. Overall survival was calculated using the Kaplan-Meier method and the impact of ARSI treatment was compared using a log-rank test with R packages Survival version 3.7 (RRID:SCR_021137)^72^ and Survminer version 0.5 (RRID:SCR_021094)^73^.

### Cistrome enrichment in accessible transposable elements

Enrichment of transcription factor cistromes in accessible elements from enriched transposable element subfamilies was performed using LOLA version 1.14 with R version 3.6.1 ^74^ and the ReMap 2022 database ^75^, with the consensus peaks overlapping all transposable elements as background. Significantly enriched cistromes were obtained (q-value <0.05 and logOddsRatio >1.5) and the top 2% recurrent cistromes in enriched transposable element subfamilies are shown. GIGGLE scores were generated using Cistromedb as described previously ^23,76^.

### Patient derived xenograft models

Paired-end ATAC-seq libraries from the LuCaP patient-derived xenograft models aligned to the concatenated mouse (mm10) and human genome (hg38) were obtained as described previously ^31^. Aligned files were processed by keeping only the properly paired reads with mapping quality of >= 30 and were deduplicated using markdup (samtools version 1.6). Reads mapping to the canonical human chromosomes were filtered and kept for peak calling as described above. TE enrichment was then performed on samples containing more than 10 million human reads and more than 15,000 peaks as described above. ChromVar was repurposed to score the transposable elements signatures compiled of uniquely enriched LINE1 subfamilies identified in the LINE1+ mCRPC subgroup.

Processed transcriptomic data from the LuCaP models was obtained from Coleman and colleagues (GEO GSE147250) ^77^. AR and NE pathway activity scores were calculated as described above.

### Visualization and statistical comparisons

Statistical analysis and data visualization using R packages were performed using the R version 4.4.2. Unless otherwise specified, data visualization was performed using ggplot2 version 3.5.1 (RRID:SCR_014601) and statistical comparisons were performed using the rstatix package version 0.7.2 (RRID:SCR_021240)^78,79^. The boxplots shown in Figure 2B, 4A, Supplementary figures 1C,E, 2A, 3B,C,E,F and 5A represent the 25th percentile, median and 75th percentile respectively. The error bars shown with the boxplots represent 1.5x the interquartile range.

## Data and code availability

The ATAC and HiC sequencing data generated from the WCDT cohort has been previously published and can be obtained from the European Genome-Phenome Archive (EGA) under the accession numbers EGAS00001006698 and EGAS00001006604 ^27,43^. The WGS has been deposited with dbGaP under the accession number phs001648. Transcriptomic and chromatin accessibility data for the LuCaP PDX models is available through (GEO GSE147250 and GSE298042 resp.). Code for data processing, analysis, and plotting is available through CodeOcean https://codeocean.com/capsule/7502073/tree/v1.

## Author contributions

L.M. and M.L. conceptualized the study. L.M. designed and conducted the experiments with help from T.M.R, G.G, A.N, S.B, T.K. K.K, S.Z.,, E.M., and M.B including the implementation of computational analyses and statistical approaches. Results were generated under the supervision of D.Q or M.L. Resources were provided by K.B, E.C, P.N, S.D, S.G.Z., W.Z, and F.F. Figures were designed by L.M. The manuscript was written by L.M. and M.L. with assistance from all authors. M.L. oversaw the study.

## Acknowledgments

The authors thank the Princess Margaret Genomics Centre and the Princess Margaret Bioinformatics group for providing support and infrastructure for the computational analysis of this work as well as high-throughput sequencing support. We thank members of the Lupien lab for their fruitful discussions and feedback. This work is supported by the Canadian Institute of Health Research (CIHR) (FRN-153234 & 168933 & 191847 to M.L.), the Canadian Epigenetics, Environment and Health Research Consortium (FRN-158225 to M.L.), the Ontario Institute for Cancer Research through funding provided by the Government of Ontario (IA 031 to M.L.) and the Princess Margaret Cancer Foundation. M.L. holds the Joey and Toby Tanenbaum Chair. Additionally, this research was supported by an SU2C-PCF Prostate Cancer WCDT Award and by the Movember Foundation. SU2C is a division of the Entertainment Industry Foundation. This research grant was administered by the American Association for Cancer Research, the scientific partner of SU2C. S.M.D, D.A.Q. and F.Y.F. were funded by the PCF Challenge Awards. S.M.D. is supported by NIH grant R01CA174777. DAQ acknowledges funding from the Benioff Initiative for Prostate Cancer Research, NCI SPORE 1P50CA275741, and Department of Defense awards W81XWH-22-1-0833 and HT94252410252. The funders had no role in study design, data collection, and analysis, the decision to publish or preparation of the manuscript. F.Y.F. and D.A.Q. were funded by a PCF Tactical Grant. The characterization and maintenance of the LuCaP PDX models was supported by the Pacific Northwest Prostate Cancer SPORE (P50CA97186), the PO1 NIH grant (PO1CA163227) and the Institute of Prostate Cancer Research (IPCR). K.H.B. is supported by NIH grants R01CA240816, R01CA276112, R01CA289390, and UG3NS132127.

## Supplementary Figure legends

**Supplementary Figure 1.**
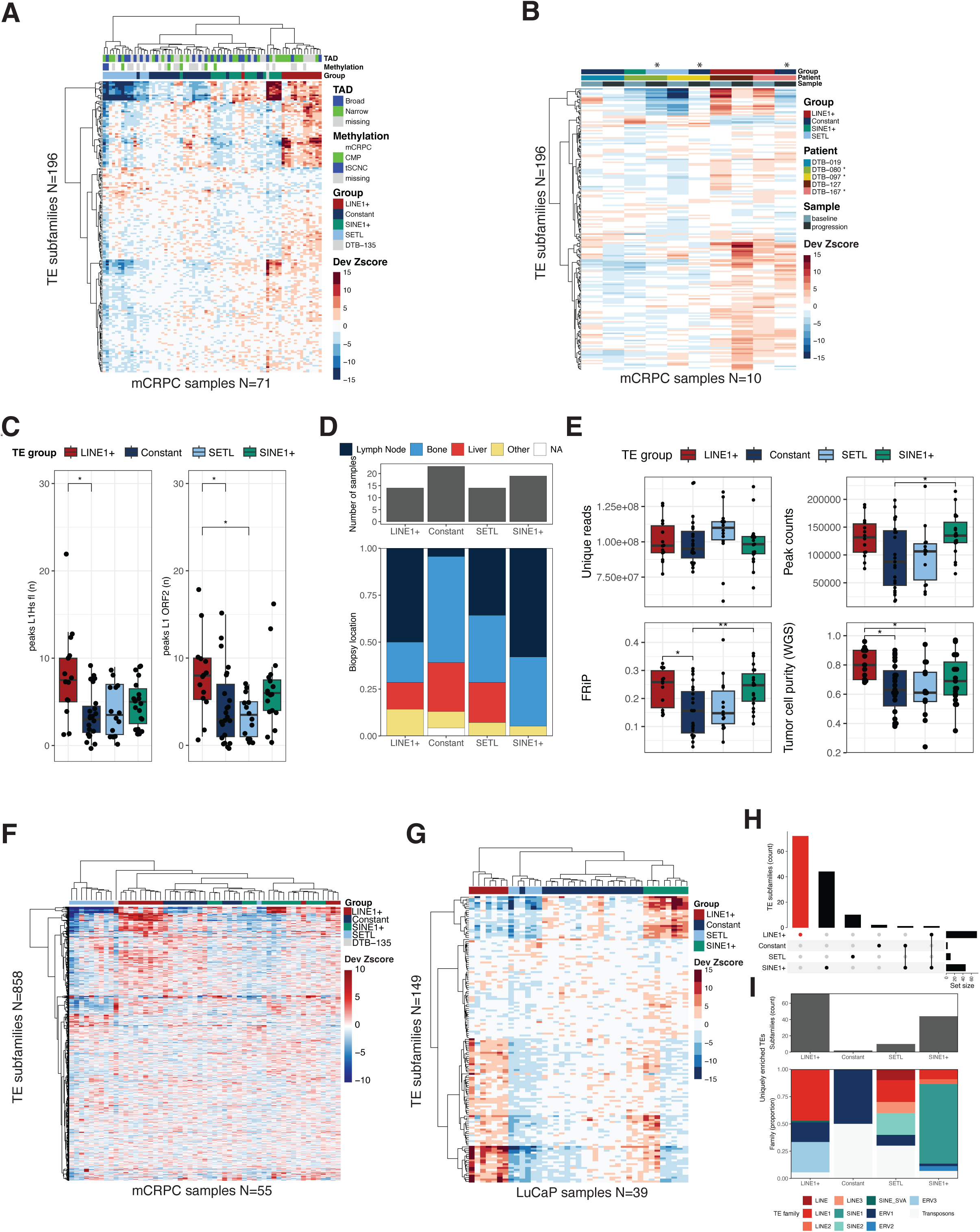
**A)** Heatmap with unsupervised clustering of deviation Z-scores (Dev Zscore) for transposable element subfamilies (N=196, rows), differentially enriched in accessible chromatin from comparing the different transposable element groups (q<0.01). The heatmap annotation (top) depicts epigenetic subtypes previously identified in the same WCDT samples. Methylation data identified the CpC hypermethylation phenotype (CMP) and treatment-induced small cell neuroendocrine carcinoma (t-SCNC) subtypes. While topology data characterized the structure of topologically associated domains (TAD) as impacting patient survival. **B)** Enrichment of transposable element subfamilies in matched samples obtained at baseline and at treatment progression from 5 patients. The heatmap shows the deviation Z-scores (Dev Zscore) for transposable element subfamilies (N=196, rows), differentially enriched in accessible chromatin from comparing the different transposable element groups (q<0.01). Samples are ordered and their annotation to the different transposable element subgroups is shown at the top. Stars indicate patients that changed transposable element subgroups upon progression. **C)** Number of peaks overlapping full-length LINE1 elements (N=107, left) and LINE1 elements with an intact ORF2 (N=146, right) by transposable element subgroup. Statistical comparison was performed using a Kruskal-Wallis test with Dunn post-hoc test and Benjamini-Hochberg corrections. L1Hs in LINE1+ vs Constant P=0.01 and ORF2 LINE1+ vs Constant and SETL P=0.02 and P=0.01 respectively. **D)** Biopsy location for the samples allocated to the different transposable element subgroups. Statistical comparison was performed by Fisher test, with pairwise comparisons and Benjamini-Hochberg corrections. For lymph node samples LINE1+, SETL and SINE1+ vs Constant P=0.006, P=0.04 and P=0.002 respectively. **E)** ATAC-seq quality metrics, including number of unique reads (top left), peaks (top right) and fraction reads in peaks (FRiP, bottom left), and tumor cell purity across the transposable element subgroups (bottom right). Tumor cell purity was estimated from whole genome sequencing. Statistical comparison was performed by Kruskal-Wallis test (number of reads, P=0.3) or an Anova (peak count, FRiP, and tumor cell content) with a Tukey post-hoc test. Peak count in Constant vs SINE1+ P=0.05, tumor cell content in LINE1+ vs Constant and SINE1+ P=0.02 and P=0.03 respectively. FRiP in Constant vs LINE1+ and SINE1+ P= 0.04 and P=0.01 respectively. **F)** Heat map with unsupervised clustering of deviation Z-scores (Dev Zscore) of transposable element subfamilies (N=858, rows) in accessible chromatin from mCRPC samples (N=55, columns) with high FRiP (>0.1) and tumor cell content (>0.5). Transposable element subgroups identified in Figure 1A are annotated. **G)** Heatmap with unsupervised clustering of deviation Z-scores (Dev Zscore) for transposable element subfamilies (N=149, rows), differentially enriched in accessible chromatin from comparing the different transposable element groups in LuCaP PDX models (q<0.05). **H)** Upset plot displaying the number of enriched transposable element subfamilies (q<0.01; median Z-score >1) shared across the different LuCaP subgroups. **I)** Barplot showing the transposable element subfamilies uniquely enriched (q<0.05; median Z-score >1) in the different LuCaP subgroups (top) and allocation to transposable element families (bottom).

**Supplementary Figure 2.**
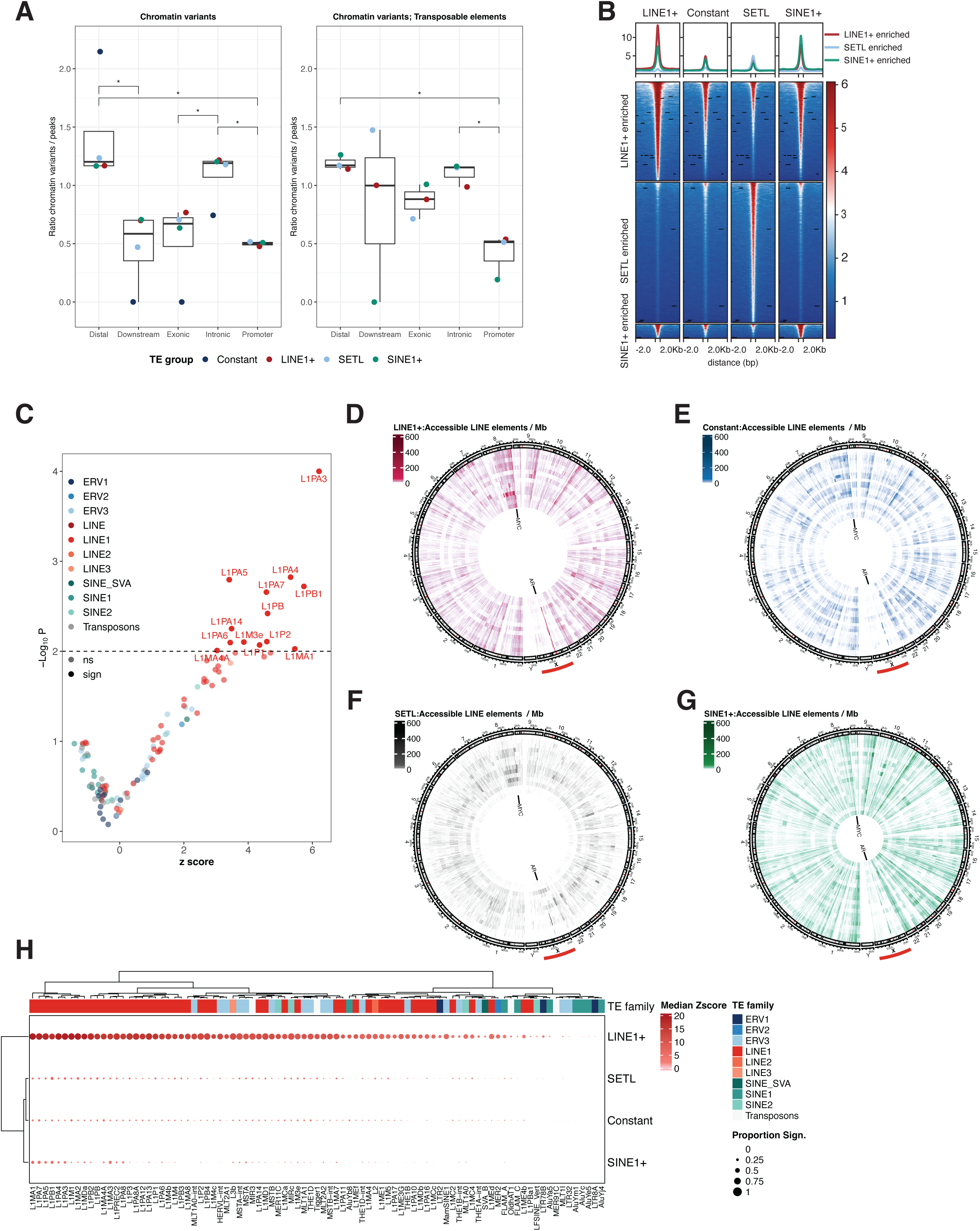
**A)** Boxplots showing the genomic distribution for chromatin variants enriched (log fold change >1, q<0.01) in the whole genome (left) or marking transposable elements from enriched subfamilies (right). The ratio of chromatin variants was normalized to the genomic distribution of all peaks and peaks marking transposable elements from enriched subfamilies. Statistical comparison was performed by t-test with Benjamini-Hochberg corrections for multiple comparisons. For all chromatin variants Distal vs Promoter and Downstream P=0.04, for Intronic vs Exonic and Promoter P=0.05 and P=0.04 respectively. For chromatin variants marking transposable elements; Distal and Intronic vs Promoter P=0.03. **B)** Tornado plot showing the average fold enriched chromatin accessibility signal in the mCRPC groups for chromatin variants identified throughout the genome (q<0.01). The length of the chromatin variants was uniformly scaled to 500 bp with 2 Kb flanking regions. **C)** Volcano plot summarizing the genomic enrichment of elements from 112 TE subfamilies within the AR amplified regions as defined by a permutation test. Significant subfamilies are labelled (P<0.01, N=13). **D-G)** Circos plot showing the genome-wide density of accessible LINE1 elements from enriched subfamilies in 1 Mb bins. Each track represents a single sample in the LINE1+ **(D)**, Constant **(E)**, SETL **(F),** and SINE1+ **(G)** groups. The position of the *AR* and *MYC* oncogenes on chrX and chr8 are annotated. **H)** Dotplot summarizing significant enrichment of accessible elements from enriched subfamilies within the *AR* amplified gene locus for LINE1+, Constant, SETL, and SINE1+ samples. Out of 112 uniquely enriched subfamilies, 89 had significant enrichment of accessible elements in the AR locus in at least one sample. The color scale represents median enrichment values (Z-score) while the size of the dot represents the proportion of samples with significant enrichment (p<0.01) in each transposable element subgroup. Unsupervised clustering was used to order the results, with the family annotation shown on top of the plotting area.

**Supplementary Figure 3.**
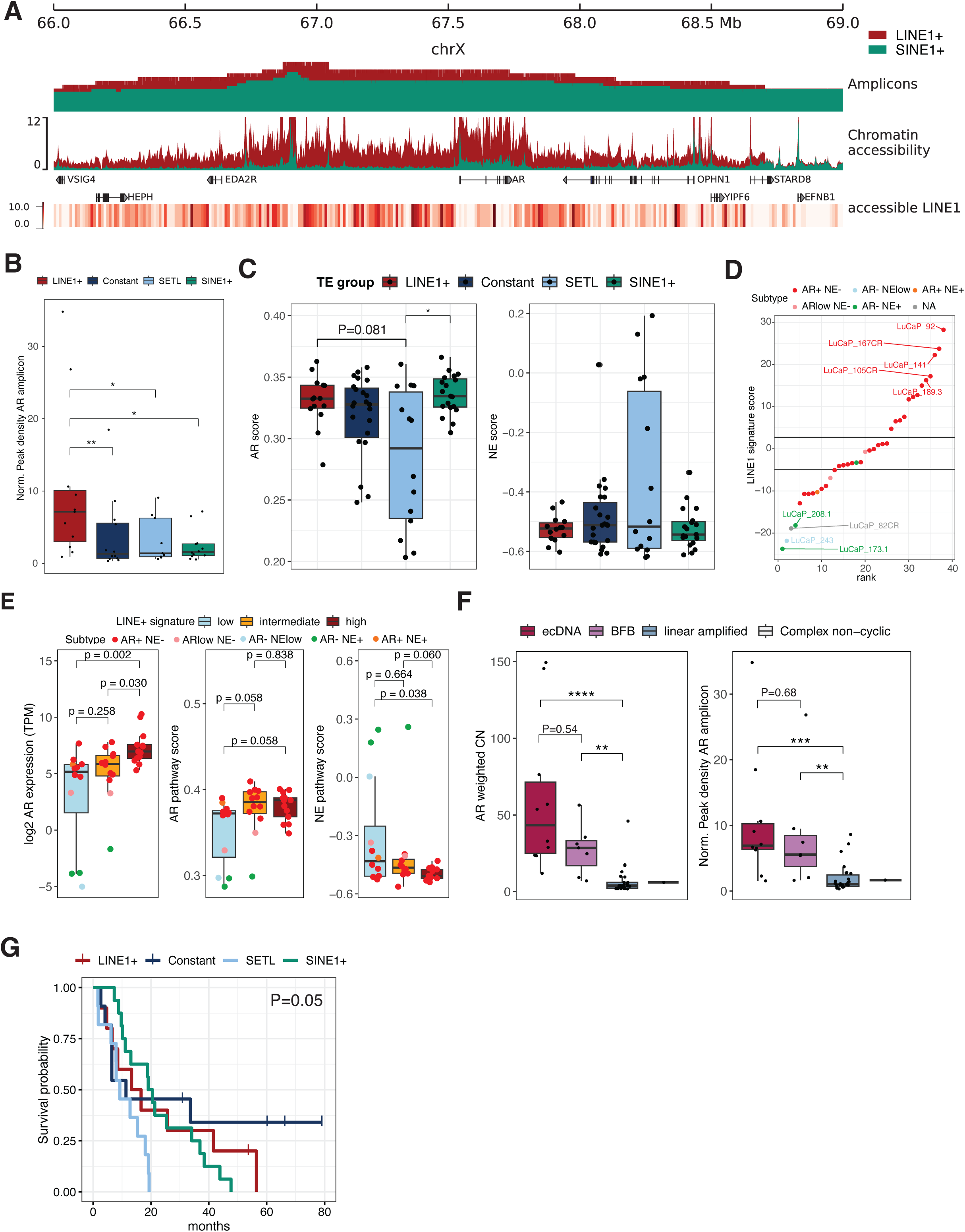
**A)** Genomic overview of the *AR* amplicons, chromatin accessibility signal, and LINE1 element enrichment for the LINE1+ (red) and SINE1+ group (green). From top to bottom; genomic annotation of the plotted region, amplicons identified in LINE1+ and SINE1+ samples scaled proportional to subgroup size, normalized chromatin accessibility signal, gene annotation, and the number of accessible LINE1 elements from enriched subfamilies in LINE1+ samples binned in 10 Kb windows. **B)** Normalized peak density for the *AR* amplicon by transposable element subgroup. Peak count was normalized to a random sampling (n=1000) of genomic regions of equal length. Statistical comparison was performed using a Kruskal-Wallis test with Dunn post-hoc test and Benjamini-Hochberg corrections. LINE1+ vs Constant, SETL and SINE1+ P=0.001, P=0.03, and P=0.03 respectively. **C)** AR and neuroendocrine (NE) pathway activity score calculated from RNA expression data. Statistical comparison across the different transposable element subgroups was performed by a Kruskal-Wallis test with Dunn post-hoc test and Benjamini-Hochberg corrections, with LINE1+ and SINE1+ vs SETL, P=0.08 and P=0.03 respectively. No significant difference was observed for the neuroendocrine score (P=0.8). **D)** Ranking the LuCaP PDX models for the mCRPC derived LINE1 signature score (Dev Zscore). The LuCaP models are annotated for their transcriptomic subtype and the horizontal lines depict the cut-offs for the top and bottom 33%. Upset plot displaying the number of enriched transposable element subfamilies (q<0.05; median Z-score >1) shared across the different LuCaP subgroups. **E)** AR expression and associated pathway activity scores in the LuCaP models scoring high (top 33%), intermediate and low (bottom 33%) for the LINE1 signature. Statistical comparison across the LINE1 signature groups was performed by a Kruskal-Wallis test with Dunn post-hoc test and Benjamini-Hochberg corrections, with AR expression and pathway activity score in low versus high LINE1 signature P=0.002 and 0.058 respectively. **F)** *AR* weighted copy number and normalized peak density by amplicon type. Statistical comparison was performed using a Kruskal-Wallis test with Dunn post-hoc corrections. Copy-numbers in *AR* ecDNA or BFB vs linear P=2.85e-05 and P=0.006 respectively. Normalized peak density in ecDNA and BFB vs linear-amplified P=5.2e-03 and P=0.009 respectively. **G)** Overall survival across transposable element groups in the WCDT cohort (N=48). Statistical comparison was performed using the log-rank test.

**Supplementary Figure 4.**
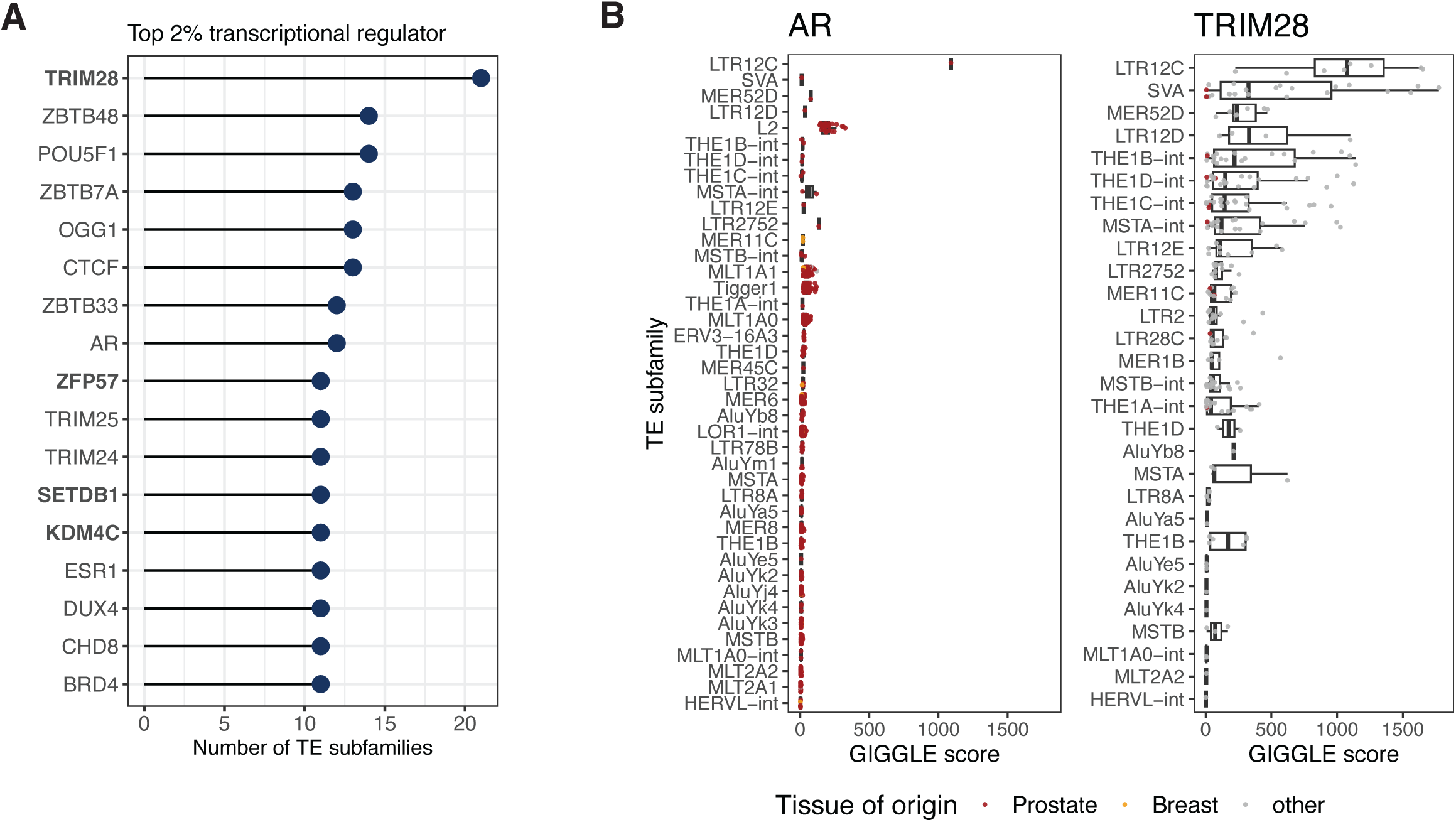
**A)** Top 2% recurring transcriptional regulators with significant enrichment (q=0.05, log-odds ratio >1.5) in nonLINE1 subfamilies (N=43) enriched in LINE1+ samples as identified from ReMaP. **B)** GIGGLE scores for AR and TRIM28 ChIP-seq data in enriched nonLINE1 subfamilies identified from the ReMaP analysis.

**Supplementary Figure 5.**
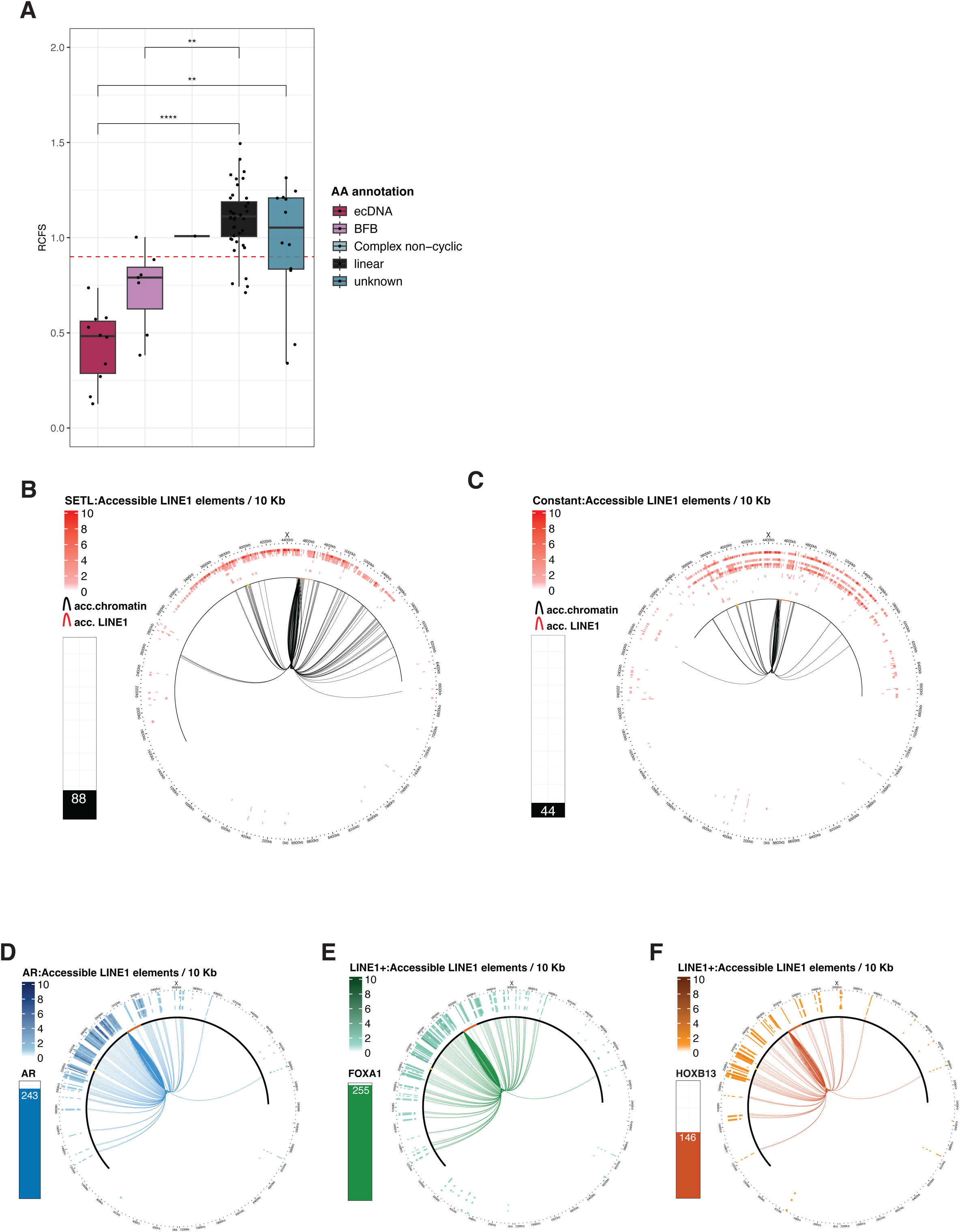
**A)** Mean scaled regional contact frequency score (RCFS) for the *AR* gene locus by amplicon status. The red dotted line indicates the RCFS cut-off (0.9) associated with AR ecDNA status. Statistical comparison was performed using a Kruskal-Wallis test with Dunn post-hoc test and Benjamini-Hochberg corrections. RCFS in ecDNA vs linear and unknown P=3.7*10^-6^ and P=0.002 respectively. RCFS in BFB vs linear P=0.007. **B-C)** Circos plots showing recurring (>25%) *AR* TSS intrachromosomal loops in SETL **(B)** and Constant **(C)** samples within chrX:63-71.8Mb (corresponding to cytoband q11.2-q13.1). Loops (inner track) anchored to accessible DNA elements with or without LINE1 sequences from enriched subfamilies are shown in red and black respectively. Barplot summarizes the number of recurrent loops shown in the Circos plot. The *AR* amplified region for the SETL and Constant subgroups is marked with a black line that includes the *AR* gene in orange and the canonical enhancers in yellow. The outer tracks show the density of accessible LINE1 elements from enriched subfamilies in 10 kb bins, for each sample. **D-F)** Circos plot showing recurring (>25%) *AR* TSS intrachromosomal loops to accessible LINE1 elements identified in LINE1+ samples, overlapping AR **(D)**, FOXA1 **(E)** and HOXB13 **(F)** binding sites. The barplot summarizes the number of recurrent loops shown in the CIRCOS plot. The AR amplicon in LINE1+ samples is marked with a black line which includes the *AR* gene in orange and the canonical enhancers in yellow. Outer tracks show the density of accessible LINE1 elements bound by AR in 10 Kb bins, for each sample in the LINE1+ group.

## Supplementary table overview

Supplementary table 1: Grouping of ATAC-seq samples from transposable element enrichment, related to Figure 1A and Supplementary Figure 1A.

Supplementary table 2: Deviation Zscores and significance values of transposable element subfamilies,, related to Supplementary Figure 1A.

Supplementary table 3: Median Z Scores of significant transposable element subfamilies, related to Figure 1B-C.

Supplementary table 4: Uniquely enriched transposable element subfamilies, related to Figure 1B-C.

Supplementary table 5: ATAC-seq quality control metrics, related to Supplementary Figure 1E.

Supplementary table 6: WGS sample ploidy and tumor cell purity, related to Supplementary Figure 1E.

Supplementary table 7: LuCaP cohort of PDX models, related to Supplementary Figure 1G-I and 2D-E

Supplementary table 8: Enrichment of accessible elements in the AR amplified locus, related to Supplementary Figure 2G.

Supplementary table 9: Copy number alterations and annotation driver genes, related to Figure 2A.

Supplementary table 10: AR normalized expression, related to Figure 2B.

Supplementary table 11: AmpliconArchitect annotation of the *AR* amplified locus, related to Figure 2D.

Supplementary table 12: Overall survival and ARSI treatment status of mCRPC patients in the WCDT cohort, related to Figure 2E-H and Supplementary Figure 3G.

Supplementary table 13: Significant transcriptional regulators enriched in LINE1 subfamilies, related to Figure 3A.

Supplementary table 14: Significant transcriptional regulators enriched in non-LINE1 subfamilies, related to Supplementary Figure 4A.

Supplementary table 15: GIGGLE scores for transcriptional regulators enriched in LINE1 subfamilies, related to Figure 3B.

Supplementary table 16: AR RCFS scores, related to Figure 4A.

